# Targeting the Annexin A1-FPR2/ALX pathway for host-directed therapy in dengue disease

**DOI:** 10.1101/2021.11.02.466887

**Authors:** Vivian V Costa, Michelle A Sugimoto, Josy Hubner, Caio S Bonilha, Celso M Queiroz-Junior, Marcela Helena Gonçalves Pereira, Jianmin Chen, Thomas Gobbetti, Gisele Olinto Libanio Rodrigues, Jordana L. Bambirra, Ingredy Passos, Carla Elizabeth Machado Lopes, Thaiane P. Moreira, Kennedy Bonjour, Rossana C. N. Melo, Milton A. P. Oliveira, Marcus Vinicius M. Andrade, Lirlândia Pires Sousa, Danielle Gloria Souza, Helton da Costa Santiago, Mauro Perretti, Mauro Martins Teixeira

**Affiliations:** Department of Morphology, Institute of Biological Sciences, Universidade Federal de Minas Gerais, Minas Gerais, Brazil; Department of Biochemistry and Immunology, Institute of Biological Sciences, Universidade Federal de Minas Gerais, Minas Gerais, Brazil; School of Medicine, Universidade Federal de Minas Gerais, Minas Gerais, Brazil; Institute of Infection, Immunity and Inflammation, College of Medical, Veterinary and Life Sciences, University of Glasgow, Glasgow, UK; William Harvey Research Institute, Barts and The London School of Medicine and Dentistry, Queen Mary University of London, London, UK; Department of Microbiology, Institute of Biological Sciences, Universidade Federal de Minas Gerais, Belo Horizonte, MG, Brazil; Department of Biology, Institute of Biological Sciences, Federal University of Juiz de Fora, Juiz de Fora, MG, Brazil; Tropical Pathology and Public Health Institute, Universidade Federal de Goiás, Goiânia, Goiás, Brazil; Department of Clinical and Toxicological Analyses, School of Pharmacy, Universidade Federal de Minas Gerais, Belo Horizonte, MG, Brazil; Centre for Inflammation and Therapeutic Innovation, Queen Mary University of London, London, UK

## Abstract

Host immune responses contribute to dengue’s pathogenesis and severity, yet the possibility that failure in endogenous inflammation resolution pathways could characterise the disease has not been contemplated. The pro-resolving protein Annexin A1 (AnxA1) is known to counterbalance overexuberant inflammation and mast cell (MC) activation. We hypothesised that inadequate AnxA1 engagement underlies the cytokine storm and vascular pathologies associated with dengue disease. Levels of AnxA1 were examined in the plasma of dengue patients and infected mice. Immunocompetent, IFNα/βR^-/-^, AnxA1^-/-^ and FPR2/ALX^-/-^ mice were infected with *Dengue virus* (DENV) and treated with the AnxA1 mimetic peptide Ac_2-26_ for analysis. Additionally, the effect of Ac_2-26_ on DENV-induced MC degranulation was assessed *in vitro* and *in vivo*. We observed that circulating levels of AnxA1 were reduced in dengue patients and DENV-infected mice. While the absence of AnxA1 or its receptor FPR2/ALX aggravated illness in infected mice, treatment with AnxA1 agonistic peptide attenuated disease manifestations. Both clinical outcomes were attributed to modulation of DENV-mediated viral load-independent MC degranulation. We have thereby identified that altered levels of the pro-resolving mediator AnxA1 are of pathological relevance in DENV infection, suggesting FPR2/ALX agonists as a therapeutic target for dengue disease.

## INTRODUCTION

Dengue is caused by one of four serotypes of dengue virus (DENV1-4) transmitted by *Aedes Aegypti* and *A. Albopictus* mosquitoes, affecting around 400 million people in 128 countries (Bhatt et al., 2013). Occasionally, dengue infection develops into a potentially lethal complication identified as severe dengue, typified by exacerbated systemic inflammation, vascular leakage, fluid accumulation, respiratory distress, severe bleeding, and/or organ impairment (Organization, 2009). No antiviral drug for dengue treatment is available, and the use of Dengvaxia^®^ (CYD-TDV), the first dengue vaccine approved by the US Food and Drug Administration (FDA), has its limitations, such as the increased risk for development of severe dengue in the immune populations (The Lancet Infectious, 2018). Thus, the combination of these factors points to dengue as a major unmet clinical problem in countries affected by this disease (Shepard, Undurraga, Halasa, & Stanaway, 2016; Stanaway et al., 2016).

The pathogenesis of severe dengue results from exacerbated host innate and adaptative immune responses to DENV. Among the target cells for DENV in humans, mast cells (MC) lining blood vessels undergo dramatic cellular activation in response to DENV, despite their high resistance to infection (Beatty et al., 2015; Modhiran et al., 2015; St John, Rathore, Raghavan, Ng, & Abraham, 2013). As essential regulators of vascular integrity, MC activation evoked by DENV triggers the production of inflammatory cytokines (cytokine storm) and vascular leakage, which ultimately result in hypovolemic shock in severe dengue (St John, Rathore, et al., 2013; Syenina, Jagaraj, Aman, Sridharan, & St John, 2015). This is substantiated by a positive correlation between circulating MCPT-1 levels and disease severity recently observed in paediatric and adult patients (A. P. S. Rathore et al., 2020; Tissera et al., 2017). In line with the suggested role of uncontrolled immune responses in the pathogenesis of DENV infection, pharmacological suppression of inflammation (Fu et al., 2014; Marques et al., 2015; Souza et al., 2009), MC stabilisation (Morrison et al., 2017), and inhibition of MC-derived protease (A. P. Rathore et al., 2019) have shown to be beneficial in experimental dengue. These studies provide proof-of-concept that host-directed therapies targeting excessive or misplaced inflammation may be a viable approach in treating severe dengue infection.

Pro-resolving mediators are cell signalling molecules synthesised in a strict temporal and spatial fashion to regulate the host response and prevent the excessive acute inflammatory reaction that damages the host (Sugimoto, Vago, Perretti, & Teixeira, 2019). The discovery of this active phase of inflammation has led to a new awareness of how a disease can emerge, including the concept that dysregulation or ‘failure’ in pro-resolving mechanisms might be involved in the pathogenesis of several chronic inflammatory disorders (Eke Gungor, Tahan, Gokahmetoglu, & Saraymen, 2014; Fredman et al., 2016; Murri et al., 2013; Schett & Neurath, 2018; Sena et al., 2013; Tabas & Glass, 2013; Thul, Labat, Temmar, Benetos, & Back, 2017; Vong et al., 2012). The pro-resolving protein AnxA1 and its cognate receptor formyl peptide receptor 2 (FPR2/ALX) are known to bear anti-inflammatory properties in sterile settings (Fredman et al., 2015; Galvao et al., 2017; Gimenes et al., 2015; Kusters et al., 2015; Leoni et al., 2015; Locatelli et al., 2014), and to exert a degree of protection in infectious settings, such as experimental tuberculosis (Tzelepis et al., 2015; Vanessa et al., 2015), sepsis (Amilcar S. Damazo et al., 2005; Gobbetti et al., 2014), pneumococcal pneumonia (Machado et al., 2020; Tavares et al., 2016) and influenza (Schloer et al., 2019). AnxA1 has been recently described to act as an endogenous modulator of MC degranulation in response to IgE/anti-IgE or compound 48/80 (Parisi, Correa, & Gil, 2019; Sinniah et al., 2019; Sinniah, Yazid, Perretti, Solito, & Flower, 2016; Yazid, Sinniah, Solito, Calder, & Flower, 2013). Since AnxA1 is well known to counter regulate overexuberant pro-inflammatory events and MC activation, we have hypothesised that an imbalance between this anti-inflammatory/pro-resolving mediator and pro-inflammatory molecules could be operating during dengue infection.

In the present work, we have analysed the role of the pro-resolving AnxA1-FPR2/ALX pathway as a regulator of excessive inflammation observed in patients with the most severe forms of dengue infection. Our results suggest that failure to trigger this molecular pathway may contribute to disease severity in dengue infection and support the AnxA1-FPR2/ALX pathway as a potential target for host-directed therapy in human dengue disease.

## METHODS

### Ethics

This study was carried out in accordance with the Brazilian Government’s ethical and animal experiments regulations (Law 11794/2008). The experimental protocol was approved by the Institutional Animal Care and Use Committee of the Universidade Federal de Minas Gerais (CEUA/UFMG, Permit Protocol Numbers 169/2016 and 234/2019). All surgeries were performed under ketamine/xylazine anaesthesia, and all efforts were made to minimise animal suffering. Human sample collection was approved by the Committee on Ethics in Research of the Universidade Federal de Minas Gerais (Protocol Numbers 24832513.4.0000.5149 and 66128617.6.0000.5149). All patients have provided signed informed consent.

### Patient recruitment

Dengue outpatients were recruited at Primary Care Center Jardim Montanhês and Santo Ivo Hospital. Inpatients were recruited in Odilon Behrens Metropolitan Hospital and Santa Casa Hospital. Healthy volunteers, negative for anti-DENV IgG (PanBio-Alere), were recruited in the community (Belo Horizonte, Minas Gerais, Brazil). Recruitment was done between the years of 2013-2016. Blood samples were obtained from 41 healthy donors and 60 dengue patients. Dengue patients were categorised into severe dengue (SD) and non-severe dengue (non-SD) groups using the 2009 World Health Organization (WHO) guidelines (WHO, 2009) and the expert physician’s judgment of disease severity (Goncalves Pereira et al., 2020). All SD patients were in-hospitalised. Of the 60 dengue patients enrolled in this study, 29 were classified as SD, and 31 were non-SD patients. Patients were included in this study if DENV infection was confirmed by dengue specific IgM capture ELISA (PanBio-Alere) and/or real-time reverse transcriptase-polymerase chain reaction (RT-PCR) conducted on all blood samples. Individuals with comorbidities such as diabetes, autoimmune diseases or obesity were excluded from this study. Serum samples collected from healthy volunteers and patients with confirmed DENV infection and a clear discharge diagnosis of either SD or non-SD were selected for measuring plasma AnxA1 levels by ELISA.

### Mice

Female BALB/c and C57BL/6 mice were obtained from the Center of Bioterism of Universidade Federal de Minas Gerais (UFMG), Brazil. Annexin A1 knockout mice (BALB/c background) (Hannon et al., 2003) and FPR2/ALX knockout mice (C57BL/6 background) (Dufton et al., 2010) were bred and maintained at animal facilities of the Immunopharmacology Laboratory of UFMG. Some experiments were conducted in type I interferon receptor-deficient mice (A129), SV129 background, obtained from Bioterio de Matrizes da Universidade de Sao Paulo (USP), bred and maintained at animal facilities of the Immunopharmacology Laboratory of the UFMG. For experiments, five-week-old wild-type mice and eight-week-old A129 mice were kept under specific pathogen-free conditions at a constant temperature (25°C) with free access to chow and water in a 12h light/dark cycle.

### Cell lines, monoclonal antibodies, and viruses

Vero and *Aedes albopictus* C6/36 cells were obtained from Banco de Células do Rio de Janeiro (BCRJ) and cultured in RPMI 1640 medium (Cultilab) or L15 medium (Cultilab), respectively, supplemented with 10% of inactivated foetal bovine serum (Cultilab). For *in vivo* and *in vitro* experiments, low passage human clinical isolates of DENV serotypes DENV-1 (EDEN 2402), DENV-2 (EDEN 3295), DENV-3 (EDEN 863), and DENV-4 (EDEN 2270) were propagated in *Aedes albopictus* C6/36 cells, and the supernatants of infected cells were harvested, filtered, concentrated, tittered by plaque assay, and stored at −80 °C until use. All *in vivo* studies with the infectious viruses were performed in a BSL-2 facility of the Immunopharmacology lab of the Institute of Biological Sciences at UFMG.

### Infections and drug treatments

For DENV infection experiments, mice received an intravenous injection of 10^6^ PFU (BALB/c WT, C57BL/6 WT, AnxA1^-/-^ and FPR2/ALX^-/-^ mice) (St John, Rathore, et al., 2013; Syenina et al., 2015) or 10^3^ PFU (A129 mice) (Costa et al., 2014; Lam et al., 2017) of DENV-2. The i.v. route of infection was chosen aiming to bypass the immune responses responsible for rapid virus clearance in natural peripheral infection (St John, Rathore, et al., 2013; Syenina et al., 2015). For treatment with the AnxA1 mimetic peptide (Ac_2-26_), BALB/c WT, C57BL/6 WT, AnxA1^-/-^ and FPR2/ALX^-/-^ mice were injected intra-peritoneally with Ac_2-26_ (150μg/animal; phosphate-buffered saline, PBS, as the vehicle) at the time of infection and daily after infection (A. S. Damazo, Yona, Flower, Perretti, & Oliani, 2006; Galvao et al., 2017; Perretti, Ahluwalia, Harris, Goulding, & Flower, 1993; Vago et al., 2012). A129 mice were treated with Ac_2-26_ (150μg/animal; i.p.) daily from day two post-infection until sacrifice (day five). Mice were randomly allocated into experimental groups using an MS Excel randomisation tool. All experiments were repeated at least twice.

### Blood parameters

Murine blood was obtained from the cava vein in heparin-containing syringes at the indicated time points under ketamine and xylazine anaesthesia (100 mg/Kg and 10 mg/Kg, respectively). The final concentration of heparin was 50 U/ml. Platelets were counted in a Neubauer chamber (Costa et al., 2014; Costa et al., 2012). Results are presented as the number of platelets per μL of blood. For haematocrit determination, blood was collected into heparinised capillary tubes (Perfecta) and centrifuged for 10 min in a haematocrit centrifuge (Fanem, São Paulo, Brazil) (Costa et al., 2014; Costa et al., 2012).

### Changes in vascular permeability

The extravasation of Evans blue dye into the liver was used as an index of increased vascular permeability, as previously described (Costa et al., 2014; Saria & Lundberg, 1983; St John, Rathore, et al., 2013). The amount of Evans blue in the tissue was obtained by comparing the extracted absorbance with a standard Evans blue curve read at 620 nm in a spectrophotometer plate reader. Results are presented as the amount of Evans blue per 100 mg of tissue.

### Cytokines, chemokines and AnxA1 quantification

The concentrations of murine CCL2, CCL5 and IL-6 in plasma samples and tissue homogenates were measured using commercially available DuoSet^®^ ELISA Development Kits (R&D). The concentrations of the MC-specific product MCPT1 in plasma samples were measured using a commercially available ELISA Ready-SET-Go!^®^ Kit (eBioscience). Human or murine AnxA1 ELISA kits (USCN Life Sciences Inc.) were used to quantify plasma levels of AnxA1. All the immunoassays were performed according to manufacturers’ instructions.

### Virus titration

A129 mice were assayed for viral titres in plasma, spleen, and liver. Blood samples were collected in heparinised tubes and centrifuged at 3000g for 15 minutes at room temperature. The plasma was collected and stored at -80 °C until assayed. For virus recovery from the spleen and liver, the organs were collected aseptically in different time points and stored at -80 °C until assayed. Tissue samples were weighed and grounded using a pestle and mortar and prepared as 10% (w/v) homogenates in RMPI 1640 medium without foetal bovine serum (FBS). Viral load in the supernatants of tissue homogenates and plasma samples were assessed by direct plaque assay using Vero cells as previously described (Costa et al., 2012). Results were measured as plaque-forming units (PFU) per 100 mg of tissue weight or per ml of plasma. The limit of detection of the assay was 100 PFU/g of tissue or per mL.

### Transaminase activity

The alanine aminotransferase (ALT) activity was measured in individual serum samples from A129 mice, using a commercially available colourimetric kit (Bioclin, Quibasa, Belo Horizonte, Brazil). Results are expressed as U/L of plasma.

### Histopathology

Liver samples from euthanised mice were obtained at the indicated time points. After that, samples were immediately fixed in 10% neutral-buffered formalin for 24 hours and embedded in paraffin. Tissue sections (4 µm thicknesses) were stained with hematoxylin and eosin (H&E) and evaluated under a microscope Axioskop 40 (Carl Zeiss, Göttingen, Germany) adapted to a digital camera (PowerShot A620, Canon, Tokyo, Japan). Histopathology score was performed as previously described (Costa et al., 2012), evaluating hepatocyte swelling, degeneration, necrosis, and haemorrhage added to a five-point score (0, absent; 1, minimal; 2, slight; 3, moderate; 4, marked; and 5, severe) in each analysis. A total of two sections for each animal were examined, and results were plotted as the mean of damage values in each mouse.

### Mast cell (MC) *in vivo* degranulation

Mice received Ac_2-26_ footpad (100μg) or i.p. (150μg) injections followed by inoculation with DENV-2 via footpad injections. Three hours later, mice were euthanised and had their hind paws removed and fixed in 10% neutral-buffered formalin for conventional histopathological processing. According to their morphological characteristics, MCs were classified as degranulated or normal, as previously described (St John et al., 2011).

### Bone marrow-derived mast cell (BMMC) generation and in vitro degranulation

BMMCs were generated as previously described (Andrade et al., 2011; Radinger, Jensen, Swindle, & Gilfillan, 2015). After four weeks, BMMCs were verified by flow cytometry to be >95% positive for the MC surface marker c-kit (CD117). To assess degranulation of BMMC in response to DENV, cells were mock-stimulated or stimulated with DENV-2 at MOI of 1, for 30 minutes, at 37 °C, in HEPES degranulation buffer (10 mM HEPES, 137 mM NaCl, 2.7 mM KCl, 0.4 mM sodium phosphate, 5.6 mM glucose, 1.8 mM calcium chloride, 1.3 mM magnesium sulphate, pH 7.4). For some groups, BMMCs were incubated with 1, 10 or 100 μM Ac_2-26_ one hour before stimulation with DENV. Degranulation was determined from the release of the granule marker β-hexosaminidase, as previously described (Andrade et al., 2011; Radinger et al., 2015). The experiment was repeated twice with BMMCs isolated from distinct animals and differentiated independently.

### Transmission electron microscopy

BMMCs pre-treated or not with Ac2-26 peptide (100 μM, 1h) followed by mock-stimulation or stimulation with DENV-2 (MOI of 1 for 30 minutes, at 37 °C) were observed by transmission electron microscopy (TEM). BMMCs obtained from three distinct animals were treated and stimulated individually. Unlike tissue-housed mast cells, primary cell cultures of the mast cells can generate cells in different maturation profiles, including early-stage and fully developed cells with typical secretory granules (Combs, 1971; JW, 1966). Thus, cells from the biological triplicate were pooled in each group to reach a suitable mature cell number in our analysis.

Following treatment and stimulation, cells were immediately fixed in a mixture of freshly prepared aldehydes (1% paraformaldehyde and 1.25 % glutaraldehyde) in 0.1 M phosphate buffer, pH 7.3, for 1h at room temperature and prepared for conventional TEM as before (Melo et al., 2009). Sections were mounted on uncoated 200-mesh copper grids (Ted Pella) before staining with lead citrate and examined using a transmission electron microscope (Tecnai Spirit G12; Thermo Fisher Scientific/FEI, Eindhoven, Netherlands) at 120 kV. A total of 203 electron micrographs were analysed to investigate morphological changes indicative of degranulation. Additionally, 1,116 secretory granules (n=372, n=360 and n=384 in the mock, DENV-2 and DENV-2+Ac_2-26_ groups, respectively) were counted in 45 electron micrographs showing the entire cell profile and the granule diameters, as well as the numbers of fused granules were quantified. Quantitative studies were performed using the Image J software (National Institutes of Health, Bethesda, MD, United States).

### RT-PCR

RNA from human plasma was obtained using PureLink Viral RNA/DNA Kits (Invitrogen). Amplifications were performed by qPCR using SuperScript III Platinum One-Step Quantitative RT-PCR System with ROX (Invitrogen) according to the manufacturer’s instructions in the presence of primers and probes described previously (Goncalves Pereira et al., 2020; Hue et al., 2011) (Supplementary Table 1). For quantifying the virus in the mouse, RNA was isolated with the RNeasy kit from Qiagen. Briefly, tissues were homogenised with the machine TissueLyserII in a small amount of the buffer using ceramic beads. Then the manufacturer’s protocol to isolate the RNA was performed, followed by cDNA synthesis. For detecting DENV-2 in the spleen, reverse primer 5’ – TTGCACCAACAGTCAATGTCTTCAGGTTC was used for cDNA synthesis, followed by RT-PCR using forward primer 5’-TCAATATGCTGAAACGCGCGAGAAACCG and reverse primer 5’-CGCCACAAGGGCCATGAACAG. For detecting DENV-2 in the liver, reverse primer 5’-GTAGCCTAGTTTGTGCAGCC was used for cDNA synthesis, and forward primer 5’-GCAGCAGAGCCATATGGT and reverse primer 5’-GTAGCCTAGTTTGTGCAGCC for RT-PCR.

### Statistics

GraphPad Prism 9.1.2 was used to determine statistical significance. Determination of sample size was based on previous publications using the software G*Power 3.1 Software. The results were analysed using appropriate statistical tests, as indicated in figure legends. Data are represented as mean ± SD.

## RESULTS

### Annexin A1 plasma levels are reduced in dengue patients

To ascertain how DENV infection impacts the expression dynamic of the pro-resolving molecule AnxA1 in humans, we measured the AnxA1 protein level in the plasma of DENV-infected patients (Fig. 1A). Dengue patients (n=60) were grouped into non-SD (dengue patients that were treated at home as outpatients, n=31) and SD (dengue inpatient cases that met WHO criteria for hospitalisation, n=29) (Goncalves Pereira et al., 2020). Demographics and laboratory characteristics are available in Table 1. Groups were comparable for sex distribution. The proportion of patients showing secondary dengue infection was not significantly different between the non-SD and SD groups. All patients with positive PCR reactions were infected with DENV-1, in line with a report that DENV-1 was the predominant serotype circulating in the city in which patients were recruited in the years of sample collection (Goncalves Pereira et al., 2020). Outpatient evolution was confirmed by remote monitoring at the convalescent phase. Interestingly, we have identified that plasma levels of AnxA1 were reduced compared to healthy controls during the acute phase of dengue infection (Fig. 1A). Stratification of groups according to disease severity at discharge showed that SD patients had discrete but significantly lower levels of AnxA1 compared to individuals with classic dengue (Fig. 1A).

**Table 1:** Demographics and laboratory characteristics of the study population from the control group, non-severe dengue (non-SD, outpatients), and severe dengue (SD, inpatients) groups during seasonal transmission 2013 to 2016.

| Characteristics and diagnosis of the study population | Control (n=41) | Non-SD (n=31) | SD (n=29) |
| --- | --- | --- | --- |
| Age <sup>a</sup> | 30 (19-58) ** | 31 (17-65) * | 42 (19-76) |
| Gender (F) | 71% (29/41) | 55% (17/31) | 52% (15/29) |
| RT-PCR (n) | 0% (0/41) | 74% (23/31) | 38% (11/29) |
| ELISA IgM (n) | 0% (0/41) | 61% (19/31) | 100% (29/29) |
| Blood collection 1-5 days after symptom onset | - | 58% (18/31) | 45% (13/29) |
| Blood collection 6-12 days after symptom onset | - | 42% (13/31) | 55% (16/29) |

<sup>a</sup> Geometric Mean (min-max). Age: Kruskal-Wallis Test P< 0.003 Gender: Fisher Test P>0.1 Blood collection: Fisher Test P>0.4

**Figure 1.**
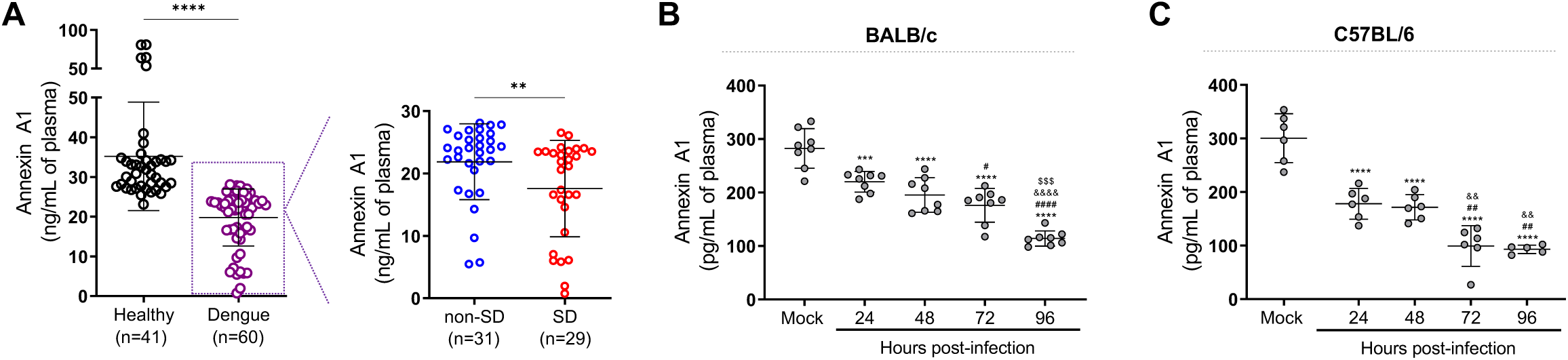
Annexin A1 levels are reduced in dengue patients and mice infected with DENV. **(A)** AnxA1 plasma levels in healthy and dengue patients. The latter group was stratified into non-severe dengue (non-SD, outpatients) and severe dengue (SD, inpatients) individuals. Each circle represents an individual participant, and horizontal bars represent mean values for AnxA1 (ng/mL of plasma), assayed by ELISA. ****p < 0.0001, **p < 0.01 by two-tailed Mann–Whitney test. **(B)** 5-week-old BALB/c (n=8) or **(C)** C57BL/6 WT (n=5-6) mice were intravenously injected with 1×10^6^ PFU of DENV-2 and culled in the indicated time points for plasma collection. AnxA1 plasma levels analysed by ELISA are shown. ***p<0.001 and ****p<0.0001 *versus* mock-infected group; ^#^p<0.05, ^##^p<0.01 and ^####^p<0.0001 *versus* 24h-infected group; ^&&^p<0.01 and ^&&&&^p<0.0001 *versus* 48h-infected group; ^$$$^p<0.001 *versus* 72h-infected group (one-way ANOVA followed by Tukey’s post hoc test). Source data 1. Raw data for Figure 1A-C.

### DENV infection reduces AnxA1 plasma levels in mice

Since we identified reduced levels of AnxA1 in the plasma of dengue patients, we sought to investigate the role of this pro-resolving mediator in dengue’s pathogenesis. To examine how the AnxA1 pro-resolving pathway operates in an immunologically intact system, we initially examined AnxA1 secretion during DENV infection over time in an immunocompetent animal model of DENV infection. Although wild type (WT) mice are more resistant to infection than immunocompromised animals (Shresta et al., 2004), these animals were proven to be productively infected by DENV and are valid hosts to investigate the mechanisms underlying DENV-induced vascular dysfunction (Chen, Hofman, Kung, Lin, & Wu-Hsieh, 2007; St John, Rathore, et al., 2013; Syenina et al., 2015). In both DENV-infected BALB/c (Fig. 1B) and C57BL/6 mice (Fig. 1C), there was a time-dependent decline in the concentration of plasma AnxA1 compared to mock-infected animals. These expression profiles motivated investigation on the role of the AnxA1-FPR2/ALX pathway in dengue disease progression and severity.

### Disruption of the AnxA1-FPR2/ALX pathway increases susceptibility to DENV infection

We then conducted experiments in different animal strains to ascertain whether changes in AnxA1 expression could impact the dynamics of DENV infection (Fig. 2A). As previously described (St John, Rathore, et al., 2013), systemic infection of WT mice caused haematological and vascular changes consistent with the human disease, systemic MC activation, and inflammatory response (Fig. 2B-D). Disease parameters were aggravated in AnxA1^-/-^ animals compared with WT mice. In the absence of AnxA1, animals showed more severe and prolonged thrombocytopenia, haemoconcentration, and vascular permeability than WT mice (Fig. 2B-D), indicating a protective role of AnxA1 in dengue disease. These findings were similar in FPR2/ALX-depleted animals (Fig. 2G-I), suggesting that these effects are due to disruption of the AnxA1-FPR2/ALX pathway. Serum levels of MCPT-1 and CCL2 were increased in either AnxA1 and FPR2/ALX KO animals compared to their respective controls after infection (Fig. 2E,F,J,K). While parameters such as haematocrit and vascular leakage returned to basal levels 72h after infection in WT animals, KO animals persisted with elevated levels, indicating a delay in resolving the host’s haematological and immune response. Interestingly, AnxA1-depleted animals have preserved their ability to control virus spread, as splenic virus replication was similar to what was found in control animals (Fig. 2-figure supplement 1).

**Figure 2.**
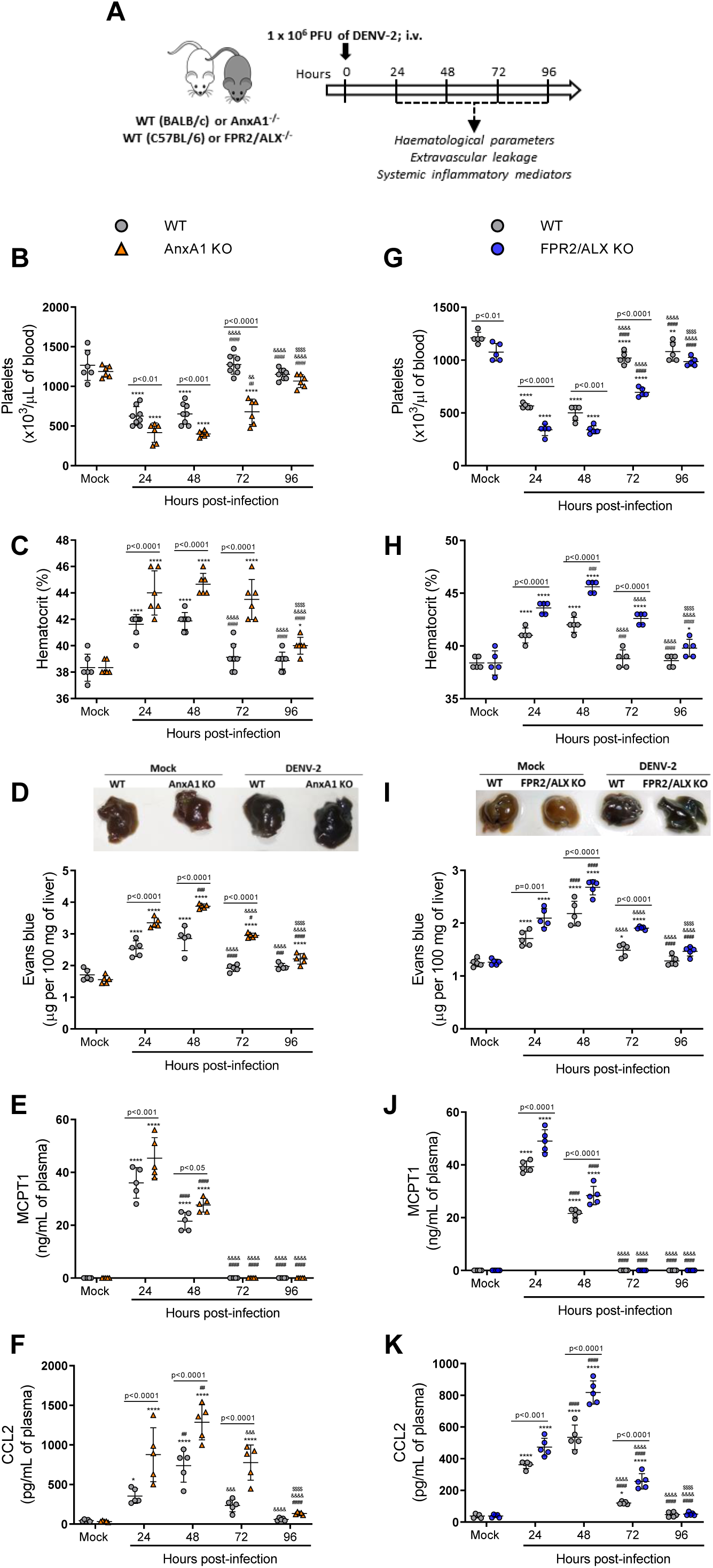
Mice are more susceptible to DENV-2 infection in the absence of Annexin A1 or its receptor FPR2/ALX. **(A)** Experimental design. **(B-F)** 5-week-old BALB/c WT and AnxA1 KO or **(G-K)** C57BL/6 and FPR2/ALX KO mice were intravenously inoculated with 1×10^6^ PFU DENV-2. Mice were culled in the indicated time points after infection, and blood and tissue were collected for the following analysis: **(B**,**G)** platelet counts, shown as the number of platelets × 10^3^/μL of blood; **(C**,**H)** haematocrit levels, shown as % volume occupied by red blood cells; **(D**,**I)** vascular leakage assay with Evans blue dye; concentrations of **(E**,**J)** MCPT1 and **(F**,**K)** CCL2 in plasma, quantified by ELISA and shown as quantity per mL of plasma. B-C, n=6-8 animals per group; D-K, n=5. Differences over time were compared by two-way ANOVA followed by Turkey’s multiple comparison test: *p<0.05, **p<0.01, ***p<0.001 and ****p<0.0001 *versus* mock-infected group; ^#^p<0.05, ^##^p<0.01, ^###^p<0.001 and ^####^p<0.0001 *versus* 24h-infected group; ^&^p<0.05, ^&&^p<0.01, ^&&&^p<0.001 and ^&&&&^p<0.0001 *versus* 48h-infected group; ^$^p<0.05, ^$$^p<0.01, ^$$$^p<0.001 and ^$$$$^p<0.0001 *versus* 72h-infected group. Differences between genotypes were compared by two-way ANOVA followed by Sidak’s multiple comparison test, as indicated in the graphs. Source data 1: Raw data for Figure 2B-K. Figure supplement 1. Supplementary DENV virus RNA quantification results.

### AnxA1 agonism attenuates DENV infection manifestations

Given the modulation of AnxA1 expression in DENV infected mice and the protective role of the AnxA1-FRP2/ALX pathway in experimental disease, we next questioned whether exogenous administration of the AnxA1 mimetic peptide Ac_2-26_ could attenuate dengue disease. Ac_2-26_ binds to FPR2/ALX and mimics most of the whole protein’s effects in experimental inflammation (Sheikh & Solito, 2018; Sugimoto, Vago, Teixeira, & Sousa, 2016). The peptide therapy in WT animals (Fig. 3A) was effective after the first injection and protected against the major changes in haematological and immune markers as evident at 24h and 48h after DENV infection compared to mock-infected animals (Fig. 3B-F). Thrombocytopenia (Fig. 3B) and haemoconcentration (Fig. 3C) were significantly milder in the group treated with Ac_2-26_. DENV-2 infection provoked increased vascular permeability in both untreated and treated animals, but to a lesser extent when animals were administrated daily with AnxA1 mimetic peptide (Fig. 3D). Of note, while increased haematocrit levels and vascular leakage were observed as early as 24h post-infection in vehicle-treated mice, treatment with AnxA1 mimetic peptide delayed the onset of both disease manifestations (Fig. 3B-C). Treatment with Ac_2-26_ also reduced the elevation in plasma MCPT-1 (Fig. 3E) and CCL2 (Fig. 3F) levels compared to the untreated group. Notably, the improvements in disease symptoms appeared to be independent of the splenic viral load (Fig. 3-figure supplement 1).

**Figure 3.**
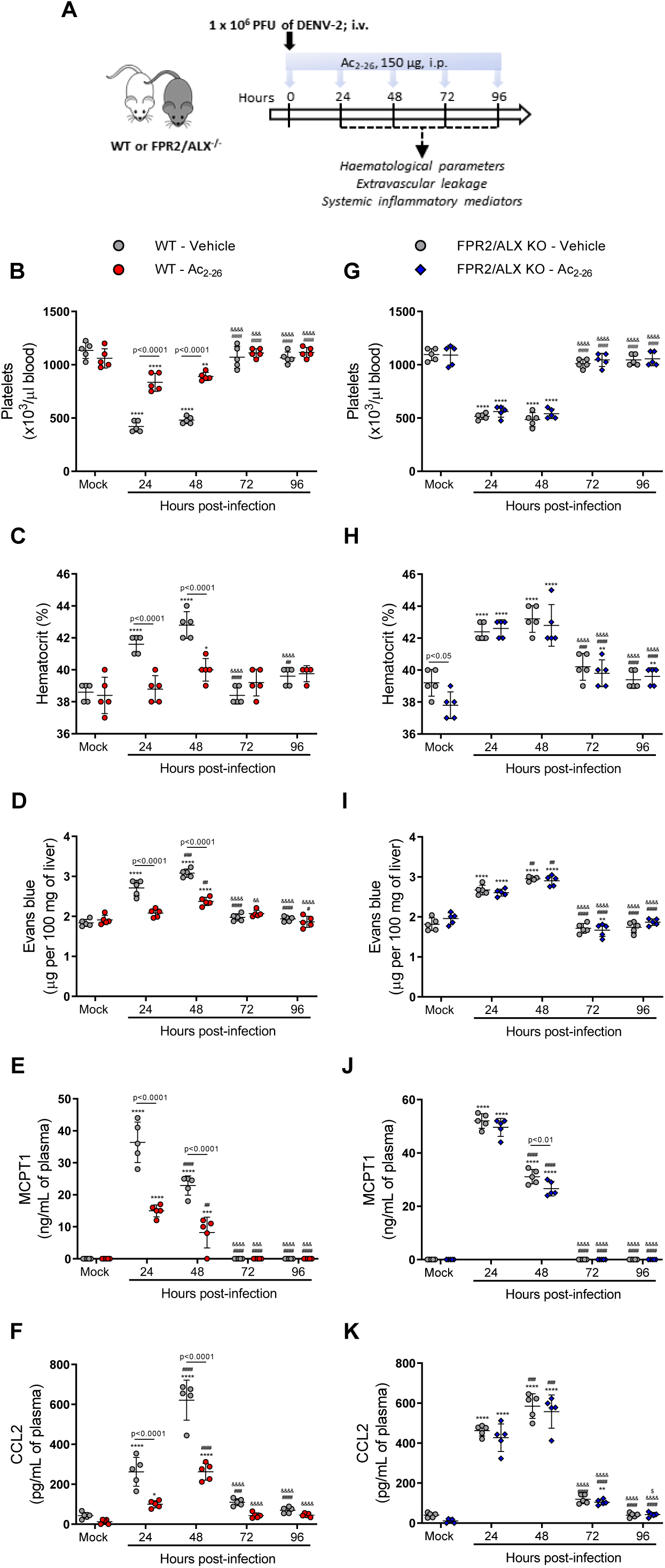
AnxA1 mimetic peptide Ac_2-26_ improves DENV-induced manifestations in WT mice and does not affect animals lacking its receptor FPR2/ALX. **(A)** Experimental design. 5-week-old BALB/c WT **(B-F)** and FPR2/ALX KO **(G-K)** mice were intravenously inoculated with 1×10^6^ PFU DENV-2. Mice were treated or not with 150μg Ac_2-26_ at the time of infection and daily thereafter by the intraperitoneal (i.p.) route. Mice were culled in the indicated time points after infection, and blood and tissue were collected for the following analysis: **(B**,**G)** platelet counts, shown as the number of platelets × 10^3^/μL of blood; **(C**,**H)** haematocrit levels, shown as % volume occupied by red blood cells; **(D**,**I)** vascular leakage assay with Evans blue dye; concentrations of **(E**,**J)** MCPT1 and **(F**,**K)** CCL2 in plasma, quantified by ELISA and shown as quantity per mL of plasma. N=5 animals per group, except for graph C where n=4-5. Differences over time were compared by two-way ANOVA followed by Turkey’s multiple comparison test: *p<0.05, **p<0.01, ***p<0.001 and ****p<0.0001 *versus* mock-infected group; ^#^p<0.05, ^##^p<0.01, ^###^p<0.001 and ^####^p<0.0001 *versus* 24h-infected group; ^&^p<0.05, ^&&^p<0.01, ^&&&^p<0.001 and ^&&&&^p<0.0001 *versus* 48h-infected group; ^$^p<0.05, ^$$^p<0.01, ^$$$^p<0.001 and ^$$$$^p<0.0001 *versus* 72h-infected group. Differences between genotypes were compared by two-way ANOVA followed by Sidak’s multiple comparison test, as indicated in the graphs. Source data 1: Raw data for Figure 3B-K. Figure supplement 1. Supplementary DENV virus RNA quantification results. Figure supplement 2. Supplementary results showing the protective effect of Ac2-26 peptide in AnxA1 KO mice challenged with DENV-2.

When we applied the same treatment schedule to FPR2/ALX KO mice infected with DENV-2, Ac_2-26_ did not affect the disease parameters under observation (Fig. 3G-K). Finally, the efficacy of Ac_2-26_ in these settings allowed us to validate the data obtained with AnxA1 KO mice, as administration of AnxA1 peptide to animals deficient in AnxA1 rescued the phenotype showed by this transgenic colony in dengue infection, bringing values of haematological and immune parameters in line with those measured in untreated mice (Fig. 3-figure supplement 2). Taken together, these results reveal the therapeutic potential of a pro-resolving peptide in the context of dengue, supporting the hypothesis that it could be operative also in settings with lower or absent AnxA1.

### Protective effects of Ac_2-26_ are independent of the control of viral loads and virus dissemination

To establish whether Ac_2-26_ treatment could have a therapeutic benefit after the infection is established and if it affects viral loads, we applied a different experimental system (Fig. 4A), using mice bearing a null mutation for the IFNα/β receptor (A129 mice). These animals are highly susceptible to DENV infection and present severe macroscopic and microscopic alterations (Costa, Fagundes, Souza, & Teixeira, 2013; Costa et al., 2014; Lam et al., 2017; Shresta et al., 2004). As seen for BALB/c and C57BL/6 strains, DENV-infected A129 mice showed reduced AnxA1 plasma levels over the course of infection, compared with mock-infected animals (Fig. 4B). While untreated A129 mice lost ∼10% of their body weight from day two post-infection until the last time point analysed, treatment with Ac_2-26_ substantially delayed weight loss onset (Fig. 4C). The AnxA1 mimetic significantly reduced thrombocytopenia and haemoconcentration in response to DENV infection (Fig. 4D,E) and displayed efficacy on controlling innate immunity mediators, with more significant effects in spleen values rather than plasma levels for CCL5 and IL-6 (Fig 4F,G). Given the characteristic of early DENV infection in our model, it is expected that the blood level of these markers would possibly not be altered by the analysed time point (5 d.p.i.). Infection of A129 mice induced a degree of liver damage, monitored by elevated serum levels of ALT and high histopathological score (Fig. 4H-J). Remarkably, treatment with Ac_2-26_ attenuated liver injury caused by DENV, as indicated by reduced histopathological score (Fig. 4H,I) and ALT transaminase levels (Fig. 4J). In this animal strain and using this protocol, we could test the effects of Ac_2-26_ following infection with other DENV serotypes and observed a significant reduction in haematological alterations, liver damage, and IL-6 production induced by either DENV-1, DENV-3, or DENV-4 (Fig. 5).

**Figure 4.**
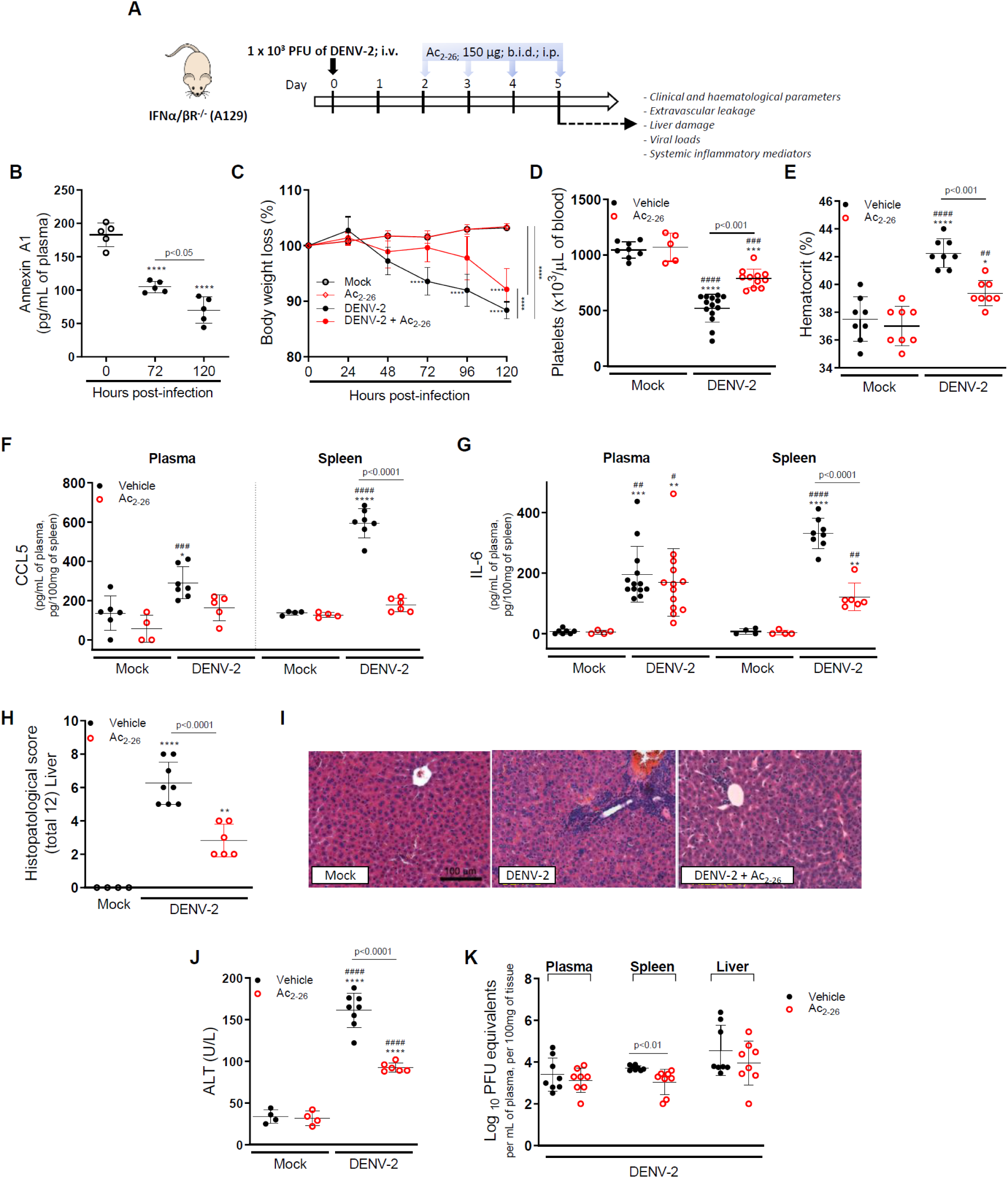
The protective effect of Ac_2-26_ administration in DENV-infected A129 mice is viral load-independent. **(A)** Experimental design. 8-week-old A129 mice were mock-infected or inoculated with 1×10^3^ PFU of DENV-2 by the intravenous (i.v.) route. From day 2, mice, were treated or not twice a day with 150μg of Ac_2-26_ by the intraperitoneal (i.p.) route. **(B)** Mice were culled in the indicated time points after infection, and plasma was collected for AnxA1 quantification by ELISA (n=5). **(C)** Bodyweight loss was assessed in the indicated time points and expressed as a percentage of initial body weight. Mock (open white circles), Ac_2-26_ (open red circles), and DENV-2-infected mice treated with vehicle (black closed circles) or Ac_2-26_ (red closed circles); n=4-8. Five days after infection, animals were culled, and blood and tissue collected for the following analysis: **(D)** platelet counts, shown as the number of platelets × 10^3^/μL of blood (n=5-14); **(E)** haematocrit levels, shown as % volume occupied by red blood cells (n=8); concentrations of **(F)** CCL5 and **(G)** IL-6 in plasma and spleen of mock and DENV-infected mice, treated or not with Ac_2-26_. Concentrations were assessed by ELISA and are shown as pg/mL of plasma or as pg/100mg of the spleen (CCL2, n=4-7; IL-6, n=4-13). **(H**,**I)** Liver of control and DENV-2-infected mice, treated or not with the AnxA1 peptide, were collected, formalin-fixed, and processed into paraffin sections. **(H)** Histopathological scores and **(I)** representative images of liver sections stained with haematoxylin and eosin. Scale Bar, 100 μm. **(J)** Plasma ALT activity represented as units/L (H-J, n=4-8). **(K)** Viral loads recovered from plasma, spleen, and liver of infected mice treated or not with Ac_2-26_, examined by plaque assay in Vero cells. Results are shown as the log of PFU/mL of plasma or as the log of PFU/mg of spleen and liver (n=8). All results are expressed as mean (horizontal bars) ± SD. In B, differences over time and between treatments were compared by one-way ANOVA followed by Tukey’s multiple comparisons test: ****p<0.0001 *versus* mock-infected animals or comparing the different groups, as indicated in the graph. In C-J, data were analysed by one-way ANOVA followed by Tukey’s multiple comparisons test: *p<0.05, **p<0.01, ***p<0.001 and ****p<0.0001 *versus* mock-infected group; ^#^p<0.05, ^##^p<0.01, ^###^p<0.001 and ^####^p<0.0001 *versus* mock-infected group treated with Ac_2-26_. In K, statistical analyses were performed by unpaired Student’s t-tests for each organ. Figure 4 – Source data 1: Raw data for Figure 4B-H,J,K.

**Figure 5.**
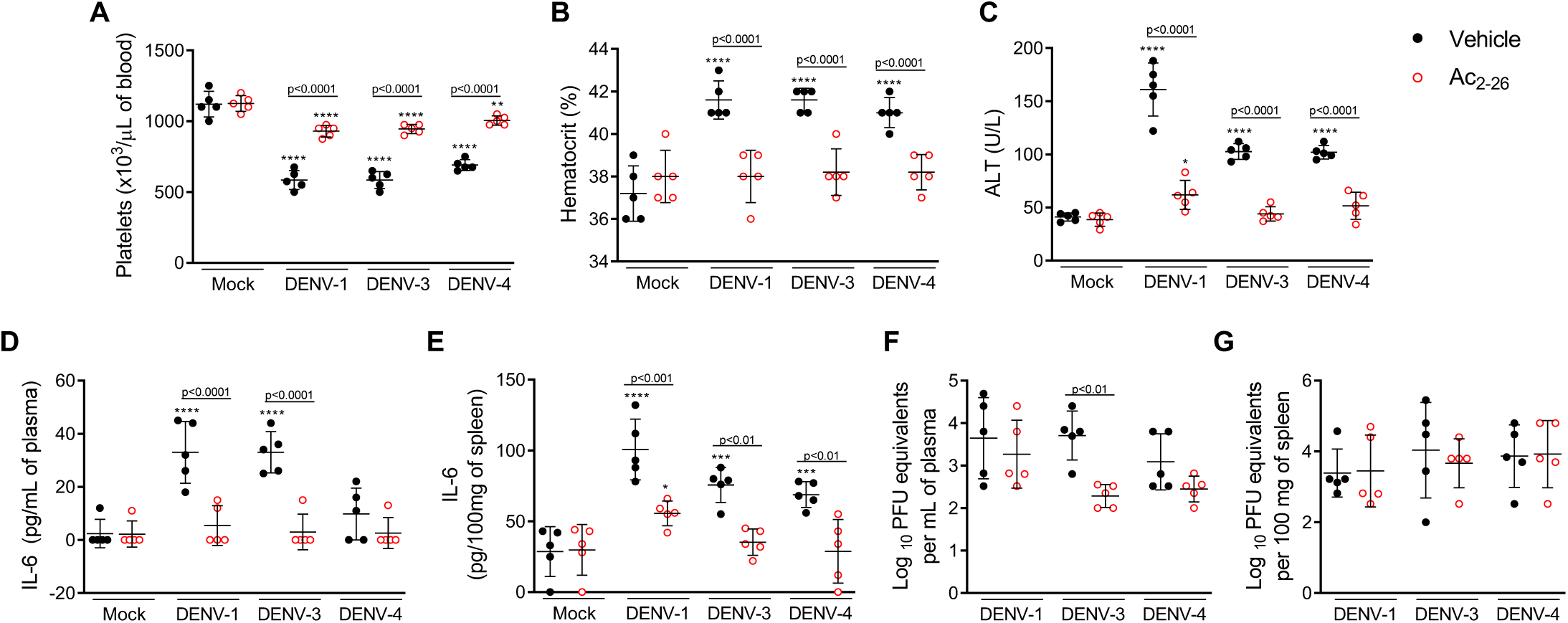
Treatment with Ac_2-26_ ameliorates disease induced by different DENV serotypes, without significantly impacting viral loads. 8-week-old A129 mice were mock-infected or inoculated with 4 × 10^4^ PFU of DENV-1, 1 × 10^3^ PFU of DENV-3 or 1 × 10^4^ PFU of DENV-4 by the intravenous route (n=5). From day 2, mice were treated with vehicle (black closed circles) or 150μg of Ac_2-26,_ i.p. twice a day (open red circles). Five days after infection, animals were culled, and blood and tissue were collected for the following analysis: **(A)** Platelet counts, shown as the number of platelets × 10^3^/μL of blood. **(B)** Haematocrit levels, shown as % volume occupied by red blood cells. **(C)** plasma ALT activity represented as units/L. Concentrations of L-6 in **(D)** plasma and **(E)** spleen of mock- and DENV-infected mice, treated or not with Ac_2-26_, assessed by ELISA. Concentrations are shown as pg/mL of plasma or as pg/100mg of the spleen. Viral loads recovered from **(F)** plasma and **(G)** spleen of mice infected with the three serotypes of DENV and treated or not with Ac_2-26_, evaluated by plaque assay in Vero cells. Results are shown as the log of PFU/mL of plasma or as the log of PFU/mg of spleen and liver. In A-E, data were analysed by two-way ANOVA followed by Dunnett’s (*p<0.05, **p<0.01, ***p<0.001 and ****p<0.0001 *versus* mock-infected group) or Šídák’s (statistical differences between infected mice treated with vehicle or Ac_2-26_, as indicated in the graphs) multiple comparison test. In F-G, statistical analysis was performed by two-way ANOVA followed by Šídák’s multiple comparison test, and differences between animals treated with vehicle or Ac_2-26_ are indicated in the graphs. Horizontal bars represent mean values. Figure 5 – Source data 1: Raw data for Figure 5A-G.

Finally, we investigated the potential impact of the Ac_2-26_ peptide on viral loads and virus dissemination. A129 mice showed systemic viral burden on day five after DENV1-4 inoculation, with detectable viremia and viral load in spleen and liver (Fig. 4K and Fig. 5F,G). Treatment with Ac_2-26_ did not affect systemic viral burden, as untreated and treated mice showed similar viremia and viral loads. Treatment with Ac_2-26_ caused only a slight reduction in viral loads in the spleen of mice infected with DENV-2 (Fig. 4K) and in the plasma of animals inoculated with DENV-3 (Fig. 5F). These data indicate that the AnxA1 mimetic positively impacts this severe dengue model, exerting little or no control on virus dissemination and viral loads, thus genuinely regulating the host response. Moreover, we show that the efficacy of the AnxA1 peptide is not restricted to a single virus serotype.

### Ac_2-26_ prevents mast cell degranulation evoked by DENV

There is compelling evidence that MC degranulation contributes to DENV-induced vascular leakage and disease severity (St John, Rathore, et al., 2013; Syenina et al., 2015; Tissera et al., 2017). In line with this, we report increased MCPT-1 levels in the plasma of DENV-infected mice (Figures 2E,J; 3E,J and Fig 3-figure supplement 2E). We hypothesised that the AnxA1 peptide could, at least in part, exert its protective effects by preventing MC degranulation. To test this hypothesis, we took advantage of *in vivo* and *in vitro* systems. We first analysed MCs from hind paw histological sections of WT mice pretreated with vehicle or AnxA1 mimetic peptide and infected with DENV (Fig. 6A). Both animals treated locally or systemically had decreased MC degranulation in comparison with control mice. To confirm a direct effect of the peptide in MC function, we cultured BMMCs and added Ac_2-26_ before challenging cells with DENV-2. We then assessed BMMC degranulation in response to DENV-2 by a standard β-hexosaminidase assay. Ac_2-26_ inhibited β-hexosaminidase release evoked by DENV-2 in a concentration-dependent manner, with ∼40% reduction in release at 100 µM Ac_2-26_ (Fig 6B). To confirm this observation, we performed a TEM assay.

**Figure 6.**
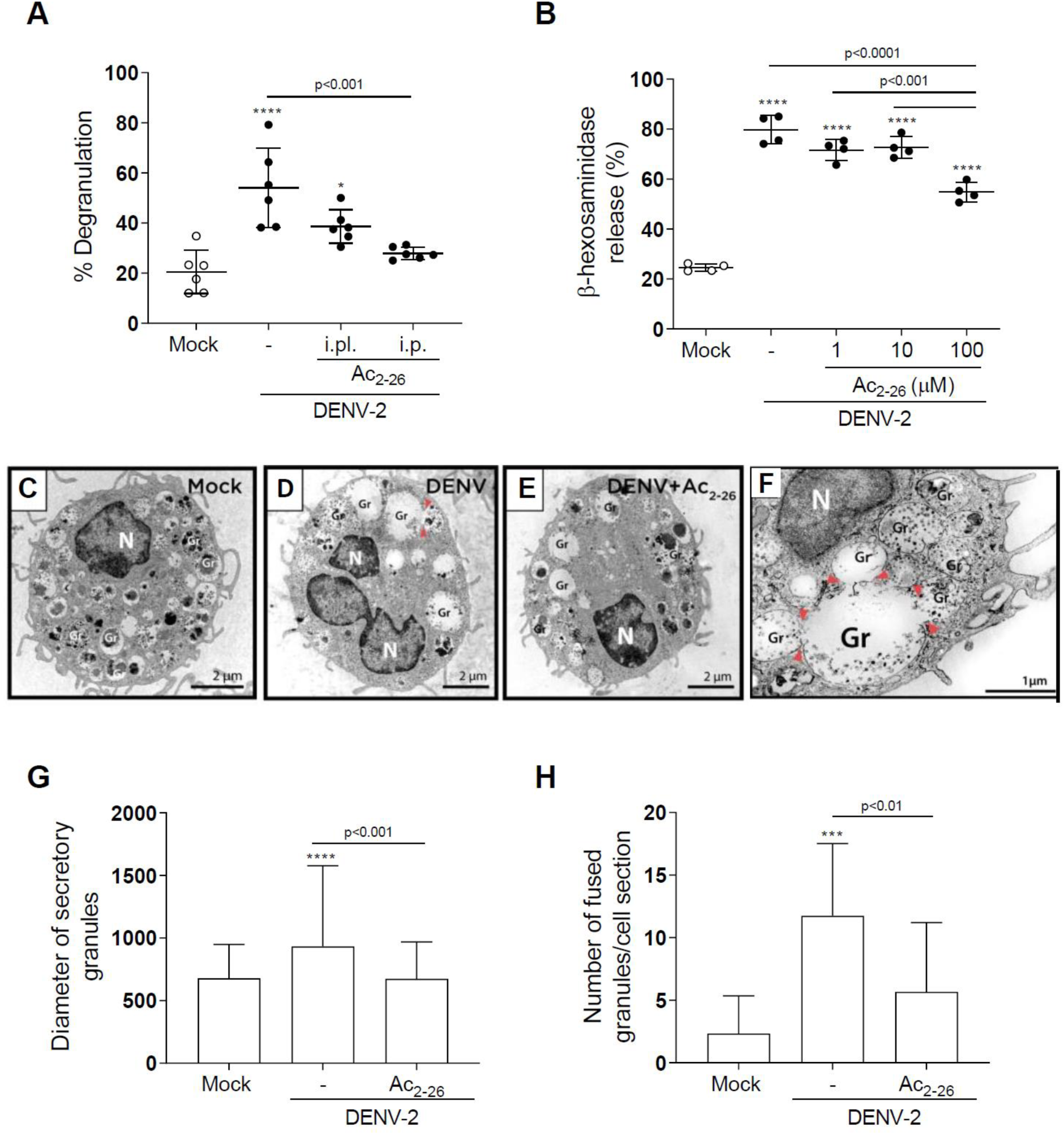
Ac_2-26_ diminishes mast cell degranulation induced by DENV both *in vivo* and *in vitro*. **(A)** WT mice were treated with Ac_2-26_ via footpad injections (i.pl.) or i.p and infected with DENV-2 via footpad injections (n=6). Three hours later, mice were euthanised and had their hind paws removed to analyse MC degranulation. **(B)** β-hexosaminidase activity of mouse bone marrow-derived mast cells (BMMC), pretreated or not with increasing concentrations of Ac_2-26_ for 1h, and stimulated with DENV-2 (MOI of 1) for an extra 30 min (n=4). The data show the percentage release of cellular β-hexosaminidase into the medium and represent two independent experiments. **(C-H)** Transmission electron microscopy (TEM) of BMMC, pretreated or not with Ac_2-26_ 100 μM for 1h and challenged with DENV-2 (MOI of 1) for an extra 30 min. **(C)** Mock-stimulated BMMC show maturing cytoplasmic granules accumulating electron-dense material. **(D)** Granule enlargement, emptying, and fusion are observed in response to DENV-2 infection. **(E)** Ac_2-26_ peptide treatment reduces morphological features of secretion evoked by DENV infection. Scale bar, 2 μm. **(F)** Granule fusions (arrowhead) in untreated DENV-infected BMMC are seen in higher magnification (scale bar, 1 μm). Significant increases in **(G)** granule diameters and **(H)** number of fused granules occur after stimulation with DENV-2 compared to both mock-stimulated cells and DENV-stimulated cells treated with Ac_2-26_ peptide. In G, bars represent the mean diameter ± SD of 372, 360, and 384 secretory granules analysed in the mock, DENV-2 and DENV-2+Ac_2-26_ groups, respectively. In H, bars represent the mean number of fused granules ± SD analysed in 15 sections per group. A,B,H, statistical analysis was performed by one-way ANOVA followed by Tukey’s multiple comparisons test: ***p<0.001 and ****p<0.0001 compared to mock-infected cells, or as depicted on the graphs. G, data analysed by Kruskal Wallis followed by Dunn multiple comparison test. Gr, secretory granules; N, nucleus. Figure 6 – Source data 1: Raw data for Figure 6A,B,G,H. Figure 6-figure supplement 1. Supplementary representative images of the footpad sections used for the quantitative analysis showed in Figure 6A.

Ultrastructural analysis of mock-stimulated cells revealed morphological features of MCs in the process of maturation with cytoplasmic granules accumulating focal, rounded aggregates of electron-dense material (Fig. 6C) (Combs, 1971; Dvorak et al., 1982). Ultrastructural evidence of degranulation was identified in BMMC cultured with DENV (Fig. 6D,F), in which cells showed enlarged cytoplasmic granules with reduced electron density and granule-granule fusion events – all morphological changes indicative of content release (Carmo et al., 2016). These features were consistently reduced when infected cells were pretreated with 100 µM Ac_2-26_ peptide (Fig. 6E). Quantification of the morphological changes in granules showed that Ac_2-26_ prevented the increase in granule diameters caused by DENV (Fig. 6G). In addition, quantitative TEM demonstrated a significant increase in the number of granule-granule fusions in response to DENV infection than the mock-stimulated group; a feature significantly attenuated upon Ac_2-26_ pre-treatment (Fig. 6H). Altogether, our data suggest that the AnxA1 mimetic peptide Ac_2-26_ directly acts on MC diminishing its degranulation in response to DENV.

## DISCUSSION

We present evidence that an inadequate engagement of the resolution circuit AnxA1-FPR2/ALX may contribute to dengue infection’s pathogenesis with particular relevance for the cohort of patients affected by the most severe forms of the disease. By exploring resolution biology as a novel approach in dengue disease, both with respect to etiopathogenesis and pharmacological opportunity, we have identified that: (i) AnxA1 is downregulated in the plasma of dengue patients in comparison to healthy individuals and in DENV-infected mice in comparison to non-infected animals; (ii) depletion of the AnxA1-FPR2/ALX pathway aggravates clinical signs and enhances MC activation associated to DENV infection, indicating a nonredundant role for this resolution pathway in the pathogenesis of dengue disease; (iii) pharmacological treatment of mice with an FPR2/ALX agonistic peptide produced beneficial effects during DENV infection; (iv) AnxA1 mimetic peptide has direct inhibitory effects on MC degranulation induced by DENV while (v) it does not seem to control viral load and virus dissemination significantly.

In recent years, a new paradigm shift has emerged in our understanding of the pathogenesis of inflammatory diseases, which results from persistent and exacerbated pro-inflammatory signals and dysregulation or ‘failure’ in resolving mechanisms (Schett & Neurath, 2018; Tabas & Glass, 2013). The severity and lethality of several infectious diseases, like dengue and the flu, frequently arise from an excessive host response, characterised by an uncontrolled release of pro-inflammatory cytokines leading to over-exuberant immune activation (Costa et al., 2013; St John, Abraham, & Gubler, 2013). Inflammation is physiologically balanced by resolution circuits, such as those centred on AnxA1 and lipid mediators (e.g. lipoxins, resolvins, protectins and maresins) that drive termination of inflammatory response yet helping the host to deal with the infective agent (Basil & Levy, 2016). In line with this, a growing body of evidence indicates that pro-resolving mediators are regulated during infection, contributing to the control and resolution of experimental and human infectious diseases (Abdulnour et al., 2016; Basil & Levy, 2016; Chiang et al., 2012; Frediani et al., 2014; Oliveira et al., 2017; Shirey et al., 2014). On the other hand, dysregulation in the production and/or action of pro-resolving mediators might contribute to the pathogenesis of sterile (Eke Gungor et al., 2014; Fredman et al., 2016; Murri et al., 2013; Sena et al., 2013; Thul et al., 2017; Vong et al., 2012) and infectious (Cilloniz et al., 2010; Colas et al., 2019; Molas et al., 2020; Morita et al., 2013) diseases, including atherosclerosis, inflammatory bowel diseases and tuberculous meningitis. Modulation of AnxA1 in the context of viral infections has been less investigated, and clinical data are scarce (Arora et al., 2016; Molas et al., 2020). Since recent evidence suggests a defective engagement of pro-resolving pathways during self-resolving infections (Cilloniz et al., 2010; Colas et al., 2019; Molas et al., 2020; Morita et al., 2013), we queried whether such alterations could also be occurring in dengue disease. Focus was given to the pro-resolving protein AnxA1 and its cognate receptor FPR2/ALX, a pathway that has been shown to exert a degree of protection in experimental tuberculosis (Tzelepis et al., 2015; Vanessa et al., 2015), sepsis (Amilcar S. Damazo et al., 2005; Gobbetti et al., 2014), pneumococcal pneumonia (Machado et al., 2020; Tavares et al., 2016) and influenza (Schloer et al., 2019). Our data indicate that in addition to the already described early induction of an inflammatory response that may be harmful instead of protective, DENV infection is also characterised by sustained downregulation of molecular components of the AnxA1 pathway. The drop in circulating AnxA1 below basal levels observed in dengue patients in this study, especially in severe dengue, suggests that the pathogenesis and severity of the disease might be associated with a failure to engage mechanisms involved in endogenous anti-inflammatory signals and its resolution, such as those centred on AnxA1. Furthermore, we have identified a protective role for the AnxA1-FPR2/ALX pathway in DENV infection, as animals presented heightened signs of disease in the absence of either ligand or receptor, compared to WT mice. In line with integrated pro-resolving properties of resolution circuits, AnxA1 and FPR2/ALX KO mice displayed diverse uncontrolled responses leading to a higher degree of susceptibility to DENV-2 infection. Markers of MC degranulation and systemic inflammation were elevated in transgenic mice compared to WT counterparts. Similarly, these genetically manipulated animals presented enhanced haematological alterations in our model of experimental dengue.

Infections are currently treated by drugs that target pathogens or inhibit their growth. In some infectious diseases, blocking inflammation pathways may be beneficial. While this approach might be successful in some infections, excessive inhibition of the immune response can also be associated with immunosuppression and increased mortality, as observed in septic patients (Carlet, Payen, & Opal, 2020). Resolution pharmacology has been proposed as an alternative to balance the host response without hampering its ability to deal with the infection. Indeed, pro-resolving receptor agonists’ exogenous administration has proven benefits in experimental infectious settings, including bacterial pneumonia (Abdulnour et al., 2016; Machado et al., 2020) and influenza (Morita et al., 2013; Ramon et al., 2014; Schloer et al., 2019). In the present work, we provide proof-of-principle that a pro-resolving tool, the Ac_2-26_ peptide, can ameliorate clinical disease and reduce circulating inflammatory mediators in an FPR2/ALX-dependent manner. The AnxA1 peptide was effective even when administered from day two after the infection onset. In addition, in our experimental models, viral load was seemingly unaffected by Ac_2-26_ treatment, supporting a protective effect independent of viral infectivity.

DENV infection initiates pro-inflammatory responses aiming to control virus spread that ultimately contributes to the immunopathology of dengue. For instance, MC activation in response to DENV plays an essential role in DENV-induced vascular pathology, particularly concerning the plasma leakage that causes hypovolemic shock in severe dengue (St John, Rathore, et al., 2013; Syenina et al., 2015). This is supported by a correlation between circulating MCPT-1 levels and disease severity in humans (A. P. S. Rathore et al., 2020; Tissera et al., 2017). It has been recently described that AnxA1 acts as an endogenous modulator of MC degranulation in response to IgE/anti-IgE or compound 48/80, suggesting that this pro-resolving axis act as a brake in MC degranulation (Sinniah et al., 2016; Yazid et al., 2013). The present study confirmed that dengue disease is associated with increased plasma levels of MCPT-1 and identified enhanced secretion in animals lacking AnxA1 or FPR2/ALX. Herein, we have identified the ability of Ac_2-26_ to reduce DENV-induced MC degranulation dose-dependently, a mechanism that might underpin the reduced MCPT-1 secretion and vascular dysfunction observed in AnxA1 peptide-treated animals. Together, *in vivo* and *in vitro* evidence suggest that Ac_2-26_, at least in part, acts by attenuating MC degranulation evoked by DENV, protecting the host against vascular dysfunction associated with the disease. This mode of action points to this pathway as a relevant potential target for DENV infection treatment, as MCs are resistant to infection but play a key role in dengue pathogenesis (Beatty et al., 2015; Modhiran et al., 2015; St John, Rathore, et al., 2013).

Our results indicate that altered levels of the pro-resolving mediator AnxA1 are of pathological relevance in dengue disease. We show that inadequate engagement of resolution circuits contributes to the excessive inflammation observed in severe DENV infection. In addition, we provide evidence for the benefits of pharmacological therapy directed to modulating host immune responses in the absence of a direct antiviral effect. These findings point to a direction for future research on applying FPR2/ALX agonists as a therapeutic target for dengue disease.

## CONTRIBUTORS

VVC, MAS, DGS, HCS, MP, and MMT conceived and planned the experiments. VVC, MAS, JH, JC, TG, GOLR, JLB, IP, CEML, and TM carried out the experiments. CMQJ performed and provided expertise for the histopathological analysis. VVC, MAS and CSB took the lead in writing the manuscript. MAS, VVC, MVMA, and JH planned and executed in vitro experiments. KB and RCNM performed, analysed, and interpreted TEM. MHGP and HCS recruited patients, performed the clinical evaluation, and collected the human samples. VVC, MAS, and LPS performed and analysed the anti-Annexin A1 ELISA assays. All authors provided critical feedback and helped shape the research, analysis, and manuscript.

## ACKNOWLEDGEMENTS

We would like to thank Ilma Marçal, Tania Colina, Frankcineia Assis, and Gilvania Santos for technical support. We also would like to thank Prof. Eng Eong Ooi from Duke NUS Medical School in Singapore for providing the DENV strains used in mouse experiments. This work received financial support from the Fapemig Hospedeiro em Dengue project, the Medical Research Council in the United Kingdom (Newton project MR/No17544/1), the National Institute of Science and Technology in Dengue and Host-microorganism Interaction (INCT dengue), a program funded by The Brazilian National Science Council (CNPq, Brazil) and Minas Gerais Foundation for Science (FAPEMIG, Brazil). JC and MP are also funded by the William Harvey Research Foundation. This study was financed in part by the Coordenação de Aperfeiçoamento de Pessoal de Nível Superior (CAPES, Brazil) – Finance Code 001. We also thank L’Oréal-UNESCO-ABC “Para Mulheres na Ciência” prize granted to VVC.

## DATA SHARING

All data has been included in the manuscript, and source data files have been provided for Figures 1-6.

**The online version of this article will include the following source code and supplementary files:**

**Figure 1 – Source data 1:** Raw data for Figure 1A-C.

**Figure 2 – Source data 1:** Raw data for Figure 2B-K.

**Figure 3 – Source data 1:** Raw data for Figure 3B-K.

**Figure 4 – Source data 1:** Raw data for Figure 4B-H,J,K.

**Figure 5 – Source data 1:** Raw data for Figure 5A-G.

**Figure 6 – Source data 1:** Raw data for Figure 6A,B,G,H.

**Figure 2-figure supplement**. Supplementary DENV virus RNA quantification results.

**Figure 3-figure supplement 1**. Supplementary DENV virus RNA quantification results.

**Figure 3-figure supplement 2**. Supplementary results showing the protective effect of Ac_2-26_ peptide in AnxA1 KO mice challenged with DENV-2.

**Figure 6-figure supplement 1**. Supplementary representative images of the footpad sections used for the quantitative analysis showed in Figure 6A.

**Supplementary Table 1:** oligo primers and probes used in clinical samples.

---

**Figure 2-figure supplement 1.**
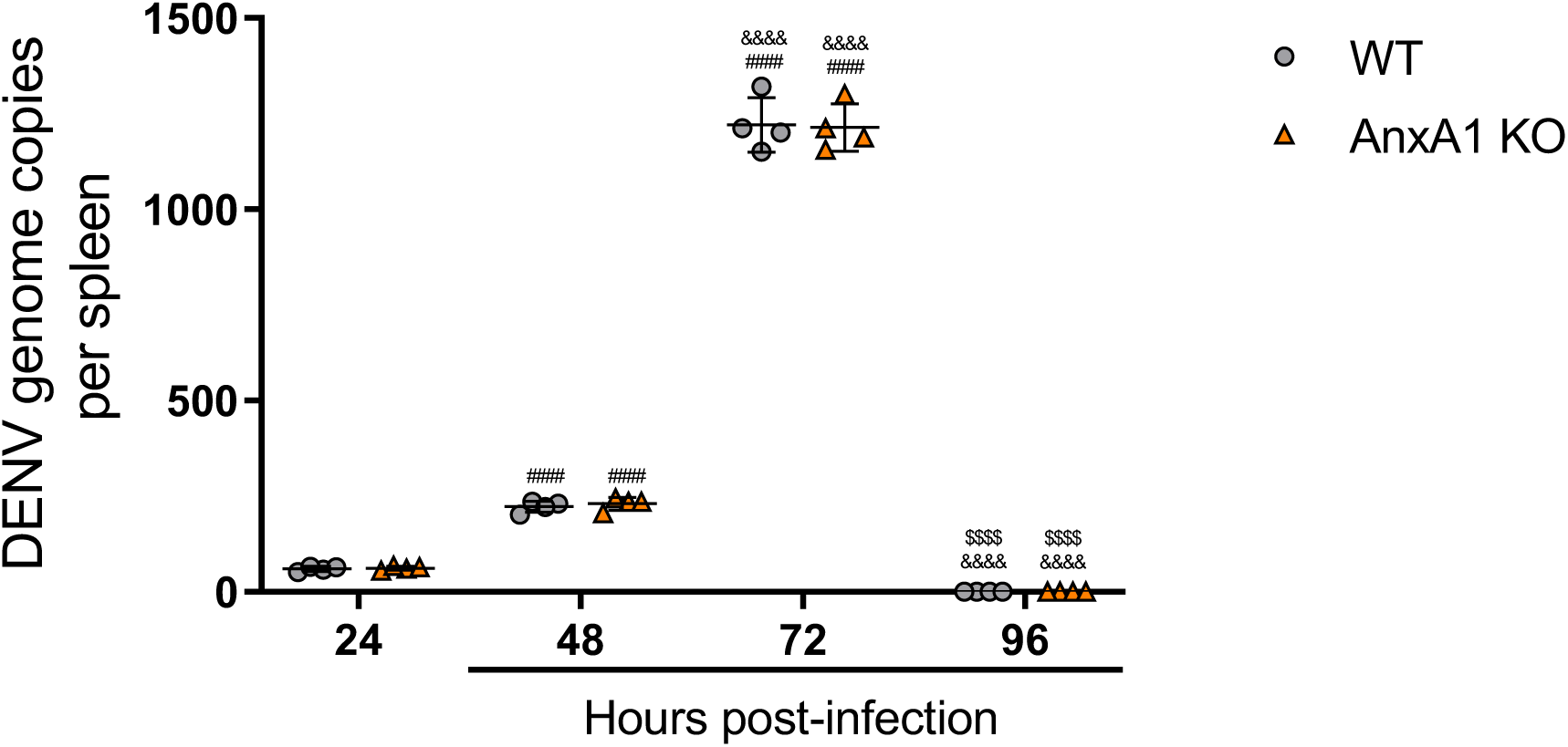
DENV virus replication is not altered by the absence of Annexin A1. 5-week-old C57BL/6 WT mice (n=4) were inoculated with 1×10^6^ PFU of DENV-2 by the intravenous (i.v.) route. Mice were culled in the indicated time points after infection, and the spleen was harvested virus RNA quantification. After cDNA conversion, real-time reverse-transcription PCR (RT-PCR) was performed to quantitate DENV genome copies in the spleen, which was normalised to spleen mass. Results are expressed as mean (horizontal bars) ± SD. Differences over time were compared by two-way ANOVA followed by Turkey’s multiple comparison test: ^####^p<0.0001 *versus* 24h-infected group; ^&&&&^p<0.0001 *versus* 48h-infected group; ^$$$$^p<0.0001 *versus* 72h-infected group. There was no statistical difference between the genotypes, as evaluated by two-way ANOVA followed by Sidak’s multiple comparison test.

**Figure 3-figure supplement 1.**
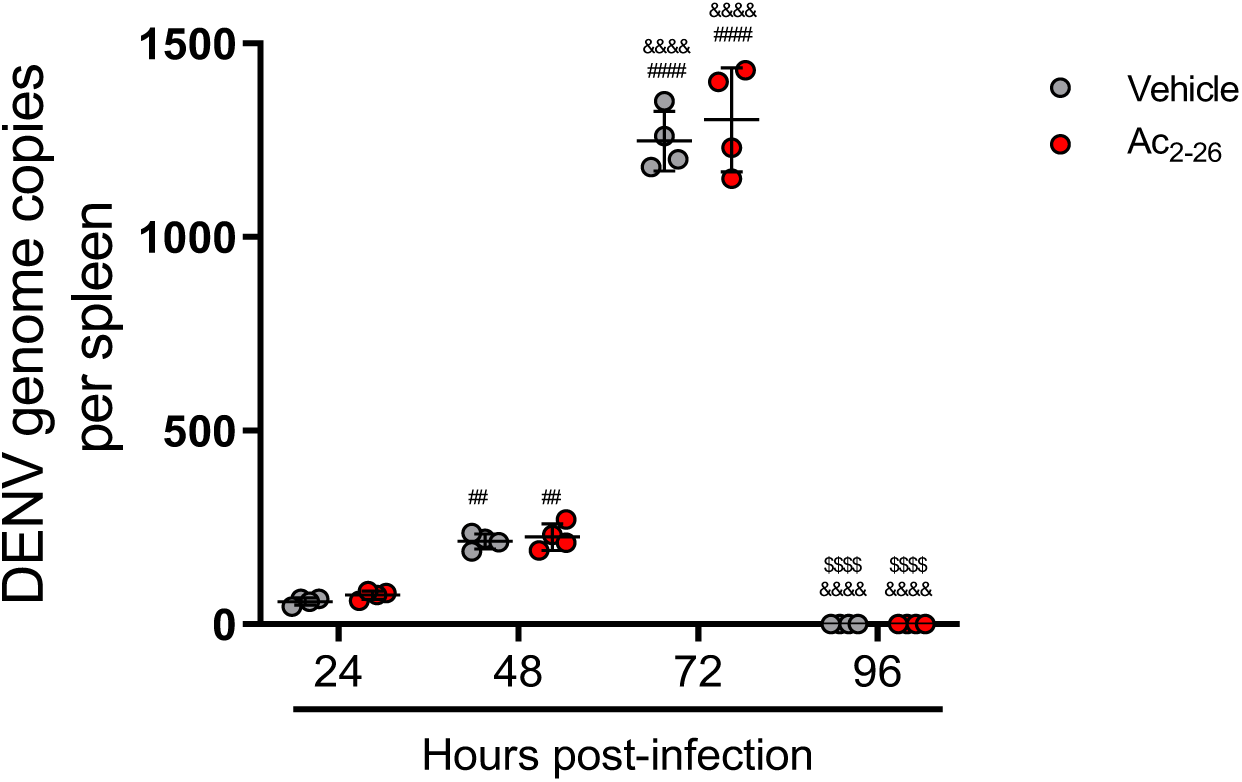
DENV virus replication is not altered by treatment with AnxA1 mimetic peptide Ac_2-26_. 5-week-old BALB/c WT mice (n=4) were inoculated with 1×10^6^ PFU of DENV-2 by the intravenous (i.v.) route. Mice were treated or not with 150μg of Ac_2-26_ at the time of infection and daily thereafter by the intraperitoneal (i.p.) route. Mice were culled in the indicated time points after infection, and the spleen was harvested virus RNA quantification. After cDNA conversion, RT-PCR was performed to quantitate DENV genome copies in the spleen, which was normalised to spleen mass. All results are expressed as mean values (horizontal bars). Results are expressed as mean (horizontal bars). Differences over time were compared by two-way ANOVA followed by Turkey’s multiple comparison test: ^##^p<0.01 and ^####^p<0.0001 *versus* 24h-infected group; ^&&&&^p<0.0001 *versus* 48h-infected group; ^$$$$^p<0.0001 *versus* 72h-infected group. There was no statistical difference between the treated and untreated groups, as evaluated by two-way ANOVA followed by Sidak’s multiple comparison test.

**Figure 3-figure supplement 2.**
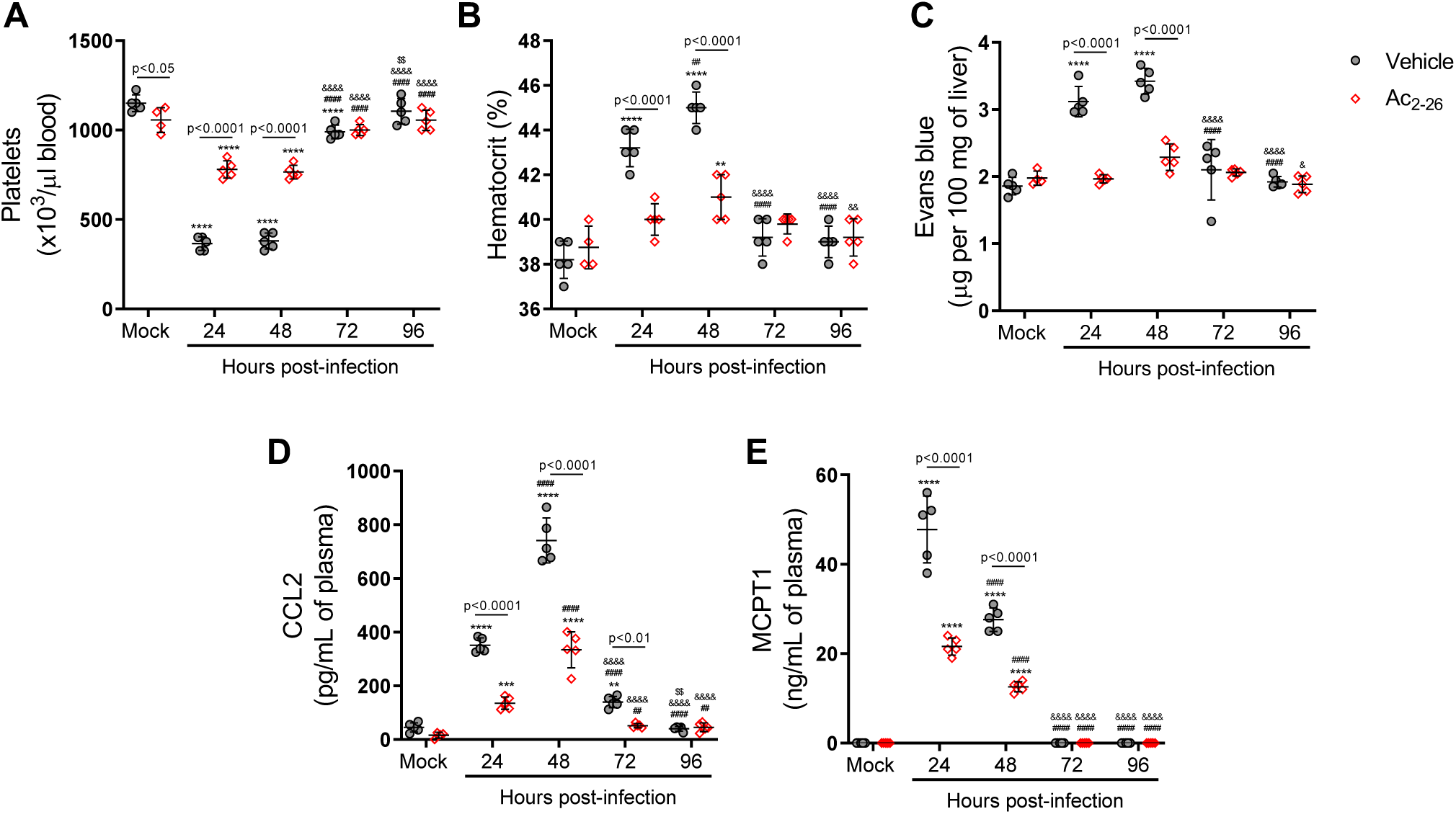
Ac_2-26_ peptide improve DENV-induced manifestations in AnxA1 KO mice. 5-week-old AnxA1 KO mice (BALB/c background) were mock-infected or inoculated with 1×10^6^ PFU of DENV-2 by the intravenous route (n=5). Mice were treated with PBS (grey circle) or 150μg of Ac_2-26_ (red open diamond) at the time of infection and daily after that by the intraperitoneal route. Mice were culled in the indicated time points after infection, and blood and tissue were collected for the following analysis: **(A)** platelet counts, shown as the number of platelets × 10^3^/μL of blood; **(B)** haematocrit levels, shown as % volume occupied by red blood cells; **(C)** vascular leakage assay with Evans’s blue dye, expressed as the amount of Evans Blue per 100mg of the liver. Concentrations of **(D)** MCPT1 and **(E)** CCL2 in plasma, quantified by ELISA and expressed as quantity per ml of plasma. All results are expressed as median (horizontal bars). All results are expressed as mean (horizontal bars). Differences over time were compared by two-way ANOVA followed by Turkey’s multiple comparison test: *p<0.05, **p<0.01, ***p<0.001 and ****p<0.0001 *versus* mock-infected group; ^#^p<0.05, ^##^p<0.01, ^###^p<0.001 and ^####^p<0.0001 *versus* 24h-infected group; ^&^p<0.05, ^&&^p<0.01, ^&&&^p<0.001 and ^&&&&^p<0.0001 *versus* 48h-infected group; ^$^p<0.05, ^$$^p<0.01, ^$$$^p<0.001 and ^$$$$^p<0.0001 *versus* 72h-infected group; or as indicated in the graphs. Differences between vehicle- and AnxA1 peptide-treated animals were compared by two-way ANOVA followed by Sidak’s multiple comparison test, as indicated in the graphs.

**Figure 6-figure supplement 1.**
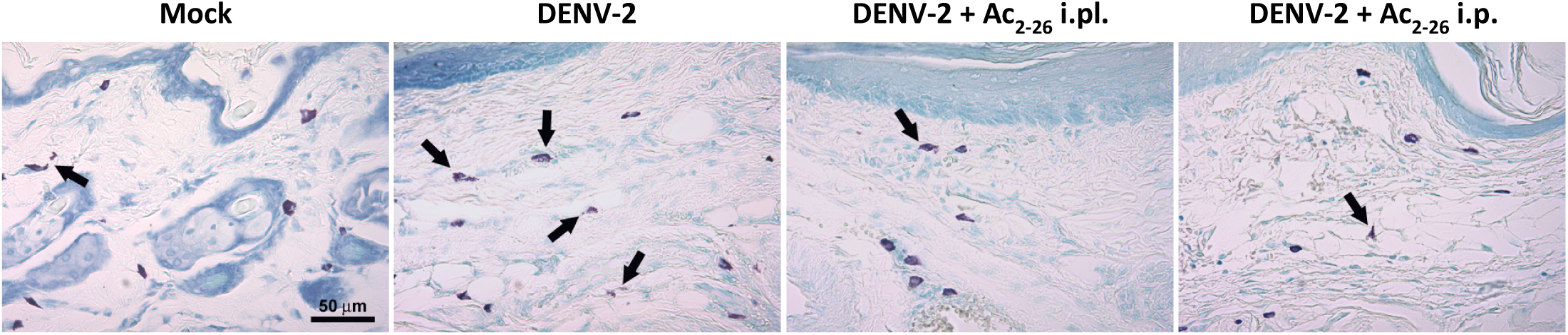
Effect of Ac_2-26_ treatment in mast cell degranulation induced by DENV-2. Representative images of footpad sections of Balb/c mice pretreated with Ac_2-26_ via footpad (i.pl.) or intraperitoneal (i.p) injections and infected with DENV-2 via footpad injections. Three hours post-infection, mice were euthanised and had their hind paws removed for the histological analysis of MC degranulation. Arrows indicate degranulated cells, visualised by toluidine blue staining.

**Supplementary Table 1:**
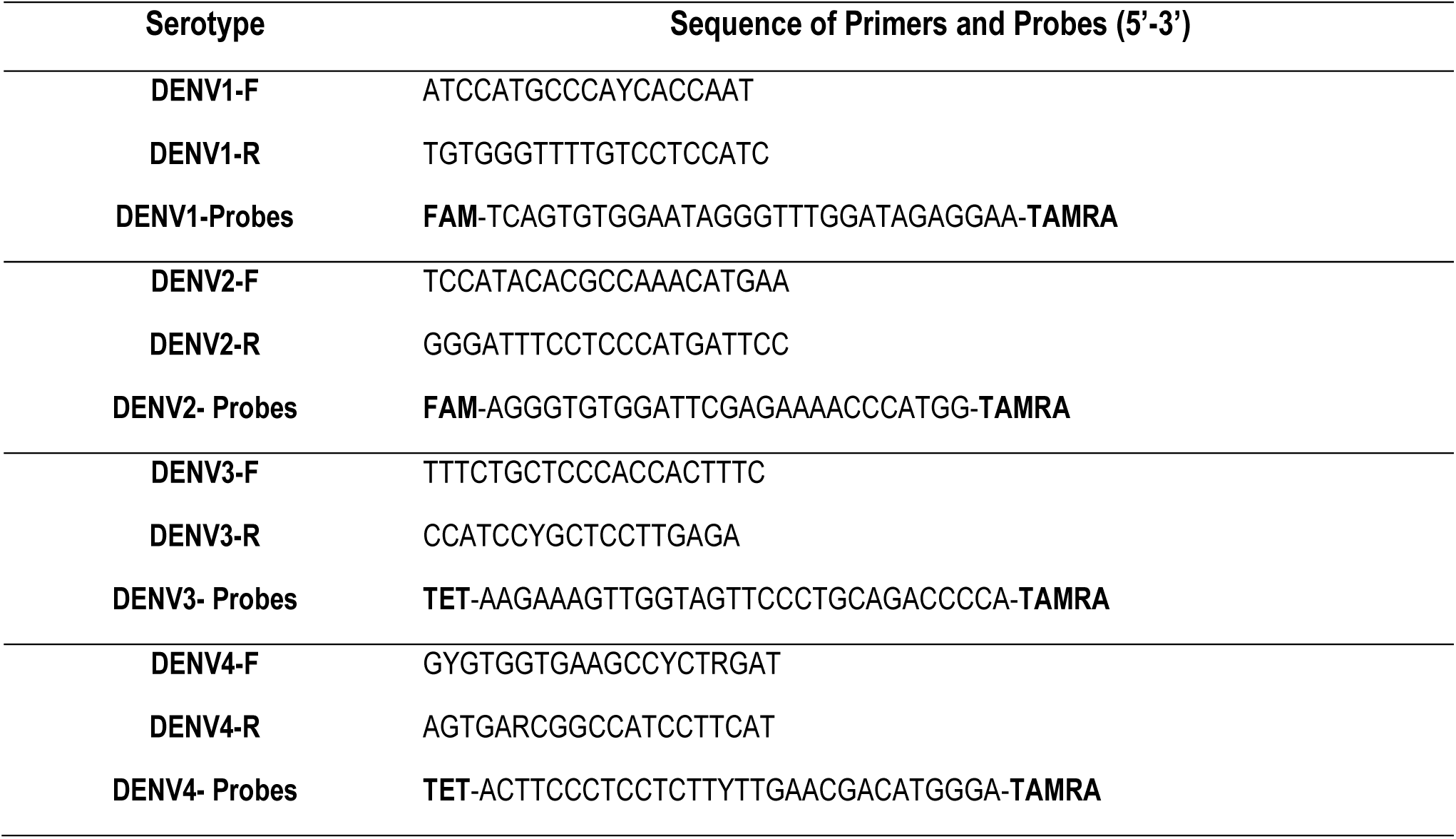
oligo primers and probes used in clinical samples.

